# A strategy to exploit surrogate sire technology in livestock breeding programs

**DOI:** 10.1101/199893

**Authors:** Paolo Gottardo, Gregor Gorjanc, Mara Battagin, R Chris Gaynor, Janez Jenko, Roger Ros-Freixedes, C Bruce A Whitelaw, Alan J Mileham, William O Herring, John M Hickey

**Affiliations:** The Roslin Institute and Royal (Dick) School of Veterinary Studies, The University of Edinburgh, Easter Bush, Midlothian, Scotland, UK; Genus PLC, 1525 River Rd., DeForest, Wisconsin 53532, USA; The Pig Improvement Company, Genus PLC, 100 Bluegrass Commons Blvd., Ste 2200, Hendersonville, Tennessee 37075, USA

**Author notes:** Email addresses: PG, GG, MB, RCG, JJ, RRF CBAW, AJM, WOH, JMH.

## Abstract

In this work, we performed simulations to develop and test a strategy for exploiting surrogate sire technology in animal breeding programs. Surrogate sire technology allows the creation of males that lack their own germline cells, but have transplanted spermatogonial stem cells from donor males. With this technology, a single elite male donor could give rise to huge numbers of progeny, potentially as much as all the production animals in a particular time period.

One hundred replicates of various scenarios were performed. Scenarios followed a common overall structure but differed in the strategy used to identify elite donors and how these donors were used in the product development part.

The results of this study showed that using surrogate sire technology would significantly increase the genetic merit of commercial sires, by as much as 6.5 to 9.2 years’ worth of genetic gain compared to a conventional breeding program. The simulations suggested that a strategy involving three stages (an initial genomic test followed by two subsequent progeny tests) was the most effective of all the strategies tested.

The use of one or a handful of elite donors to generate the production animals would be very different to current practice. While the results demonstrate the great potential of surrogate sire technology there are considerable risks but also other opportunities. Practical implementation of surrogate sire technology would need to account for these.

## Introduction

In this study, we performed simulations to develop a strategy for exploiting surrogate sire technology [1-2] in animal breeding programs (Fig. 1). Surrogate sire technology allows the creation of males that lack their own germline cells, but have transplanted spermatogonial stem cells from other donor males. The concept requires the production of recipient males with an ablated germ line. Rodent males can have their germline ablated using chemotoxic drugs or localised irradiation of the testes, but, importantly for use in livestock breeding, this ablation is incomplete and recipient sperm output is mixture of donor and recipient cells [3]. The mammalian NANOS2 gene seems to be absolutely required for the maintenance of germ line cells in males only [4]. In mice, Nanos 2 knock out males the testes completely lack germ-line cells, but there is no effect in females [4]. NANOS 2 knock out pigs have been produced using CRISP/Cas9 gene editing [1] and boars homozygous for the knockout likely provide ideal recipients for the surrogate sire concept.”

**Fig. 1.**
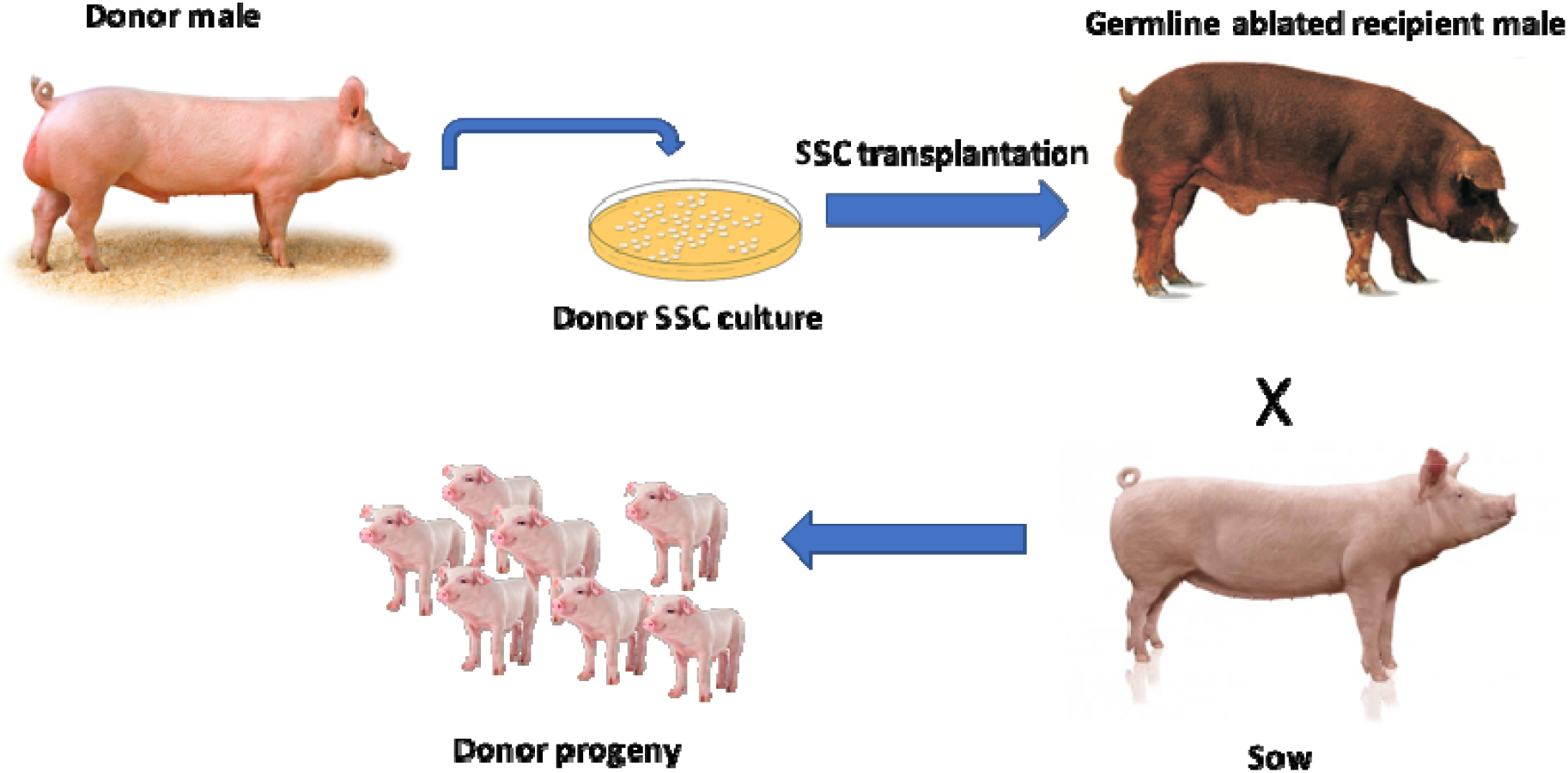
Schematic depicting the possible application of spermatogonial stem cell transplantation methodology in pig production (depiction inspired by Oatley et al., 2018 [2])

With this technology, a single elite male donor could give rise to huge numbers of progeny, potentially as much as all the production animals in a particular time period. This potential offers many advantages. Firstly, it would reduce the genetic lag between the elite nucleus animals and the production animals. Secondly, it could enable better matching of specific management plans to the genetics. Thirdly, as we outline in the discussion it could enable exploitation of combining ability. The latter could increase production on farm and increase investment and innovation in breeding by enabling a greater ability to protect intellectual property.

Typically, animal breeding programs are implicitly or explicitly organized in pyramid structures with layers (Fig. 2). The top layer is the nucleus, which is improved using recurrent selection. Nowadays most selection decisions are made using genomic based testing rather than traditional phenotype based testing [4–8]. The middle layer is the multiplication, where the nucleus genetics is multiplied and sometimes crosses between purebred lines are produced. The base layer is the commercial sector, where the majority of animals are kept for production. The commercial producers often make a final cross between the terminal line sires and the maternal line dams.

**Fig. 2.**
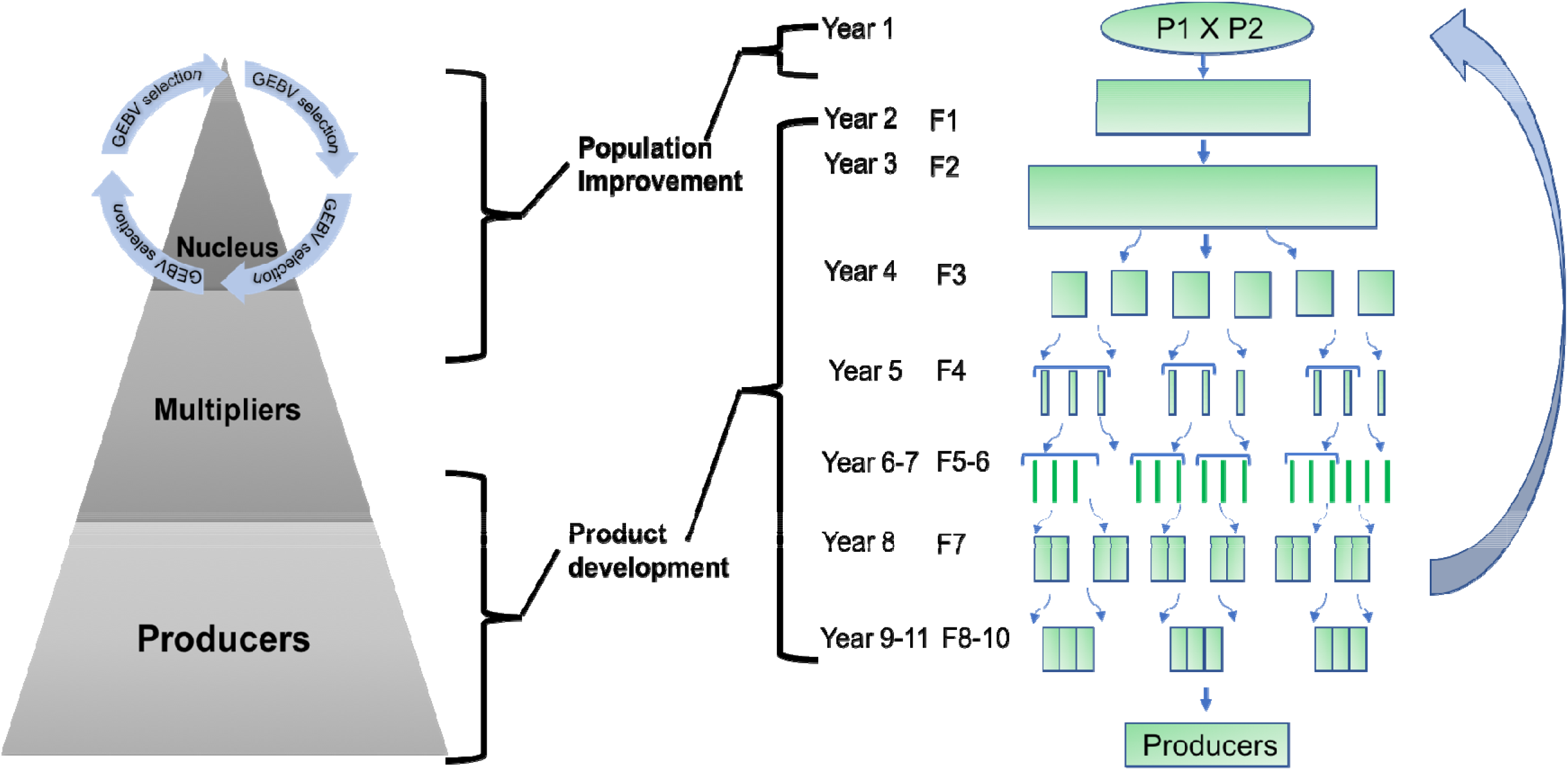
Example animal (left) and plant (right) breeding schemes

The need to generate huge numbers of production animals and the limited number of progeny that a male can produce means that large numbers of nucleus animals must contribute genetics to the subsequent layers and that one to several generations are required for multiplication. These factors give rise to a genetic lag, a difference in genetic mean between the nucleus and commercial layers. This lag can also be represented with the number of years of genetic gain [9], e.g., ~4 years in a pig breeding program. Surrogate sire technology would allow a single elite nucleus male to give rise to very large numbers of commercial animals, by donating spermatogonial stem cells to its commercial surrogates [1]. This could shorten the lag between the nucleus, multiplication, and commercial layers.

Using surrogate sire technology in this way would require that animal breeding programs identify elite donor males and create surrogate sires. This process should take place in a sufficiently small amount of time so that the extra genetic gain would not be significantly reduced by the extra time required for the identification of donors and creation of surrogate sires.

A restructured animal breeding program with surrogate sire technology would be conceptually similar to a plant breeding program that produces clonally propagated individual lines or inbred lines or hybrid lines (Fig. 2). These programs seek: (i) to identify the best individual (note: here we take individual to mean clonal, inbred or hybrid lines), or a handful of individuals, from a population of individuals; and (ii) to disseminate this individual very widely in the commercial layer [10]. To identify the best individual, plant breeders typically use multiple stage testing and selection. As the breeding program progresses through these stages the number of individuals being tested is reduced and the precision of these tests increases. The small number of individuals in the final stages are intensively tested in large replicated experiments that are repeated across several environments and years. This ensures that the commercially released individual is well characterized and carries a minimal risk of major undetected weakness. This is necessary because this individual will have a huge footprint in the commercial layer. Similar levels of evaluation would be needed with surrogate sire technology in animal breeding programs.

The objective of this study was to develop a strategy for exploiting surrogate sire technology in animal breeding programs. This strategy involved a subtle, but important, reorganisation to combine components of traditional animal and plant breeding programs. The reorganization is similar to the two-part breeding program that we recently proposed for the incorporation of genomic selection into plant breeding programs [11]. The reorganization involves an explicit partitioning of a breeding program into a population improvement component and a product development component. The population improvement component is similar to the currently used recurrent genomic selection in many animal breeding nucleus populations. The product development component is similar to traditional plant breeding programs and involves a number of stages of testing to identify the elite donors. The product development component could make use of testing for combining ability, if that was appropriate for the particular species of interest.

With a focus on application in pig breeding, several alternative versions of the reorganized breeding program were compared to different variants of a conventional breeding program using simulation. The alternative versions varied: (i) the number of stages of testing; (ii) the number of donor candidates tested at subsequent stages; (iii) the accuracy of the genomic test at the first stage; and (iv) the accuracy of progeny test in later stages. The results showed that using surrogate sire technology would significantly increase the genetic merit of commercial sires, by as much as between 6.5 and 9.20 years’ worth of genetic gain compared to different variants of a conventional breeding program. The simulations suggested that an identification strategy involving three stages (a genomic test followed by two subsequent progeny tests) was the most effective of all the strategies tested. The use of one or a handful of elite donors to generate the production animals would be very different to current practice. While the results demonstrate the great potential of surrogate sire technology there are considerable risks and these are discussed.

## Methods

Simulation was used to evaluate the impact of a strategy for exploiting surrogate sire technology in animal breeding programs. One hundred replicates of various scenarios were performed. Scenarios followed a common overall structure but differed in the strategy used to identify elite donors and how these donors were used (Fig. 3, 4).

**Fig. 3.**
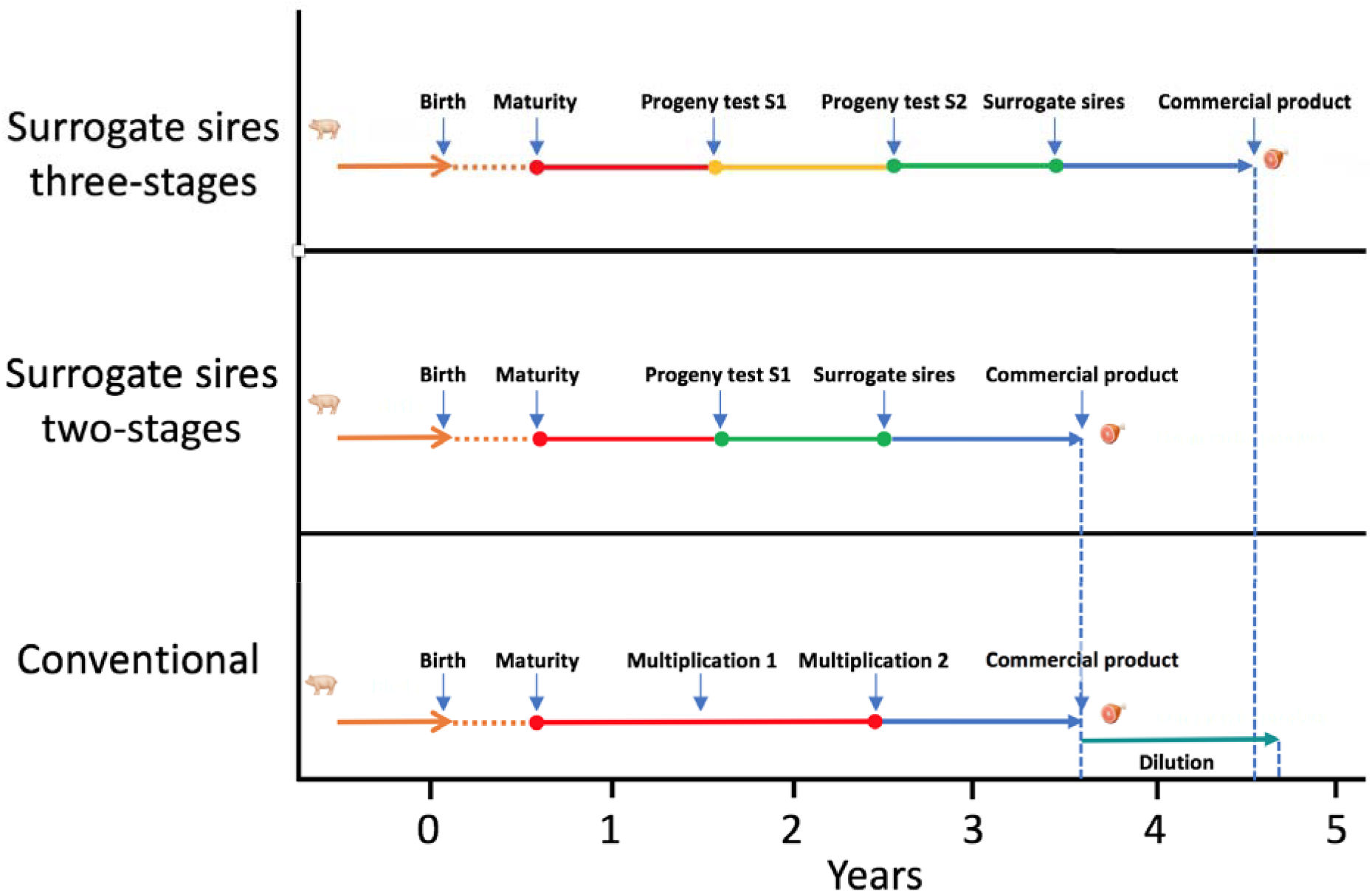
Timeline of the different strategies to identify and disseminate genetic improvement

**Fig. 4.**
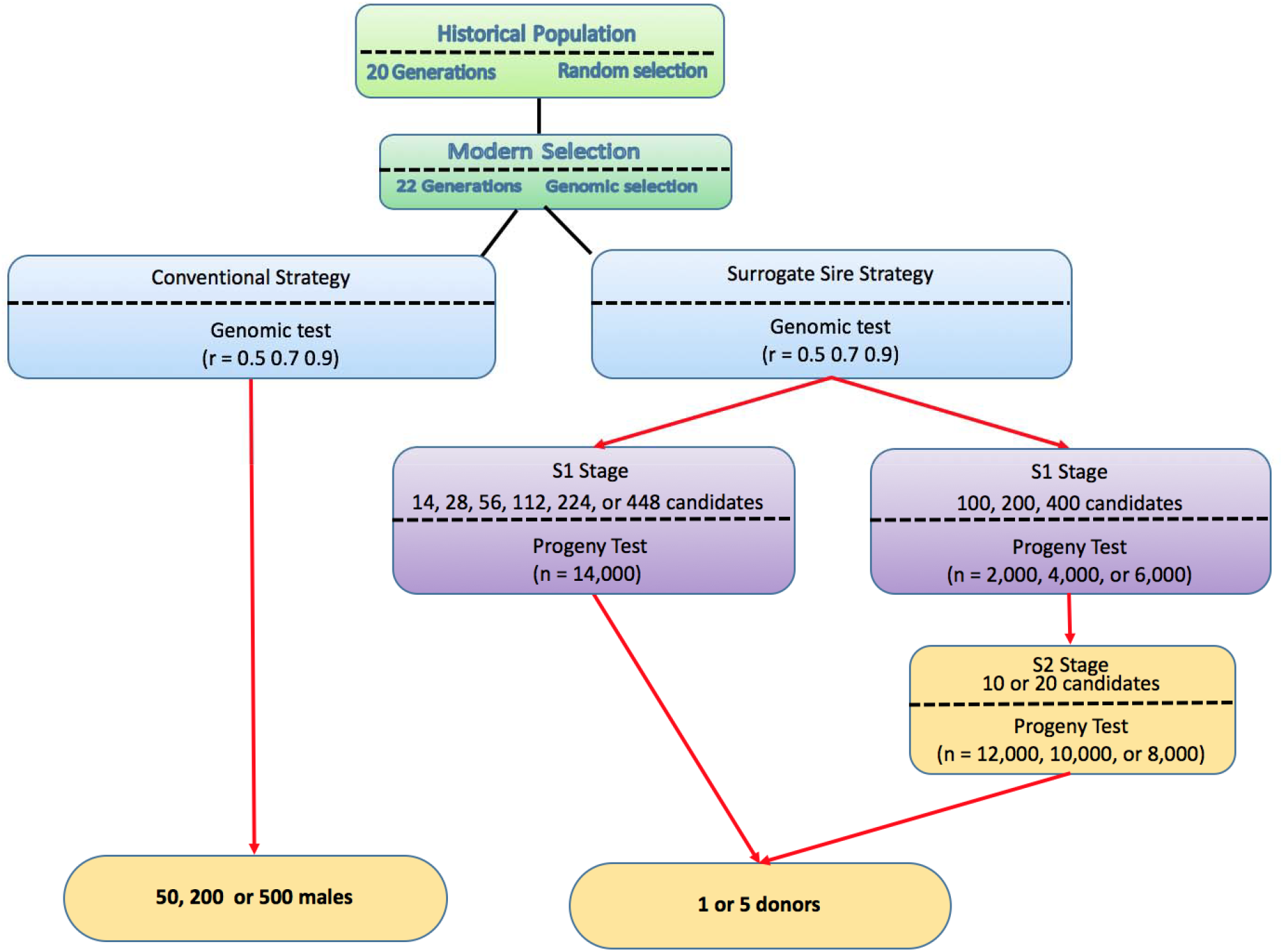
Map of the scenarios used in the study

Conceptually, the simulation scheme was divided into historical and future phases. The historical phase represented historical evolution and recent animal breeding efforts up to the present day, under the assumption that animal populations have evolved for tens of thousands of years, followed by 22 recent generations of modern animal breeding with selection on genomic breeding values in a nucleus population. The future phase represented 20 future generations of modern animal breeding, with selection on genomic breeding values in a nucleus population that subsequently supplied genetic improvement to multiplication and commercial layers. The historical animal breeding generations were denoted −21 to 0 and the future animal breeding generations were denoted 1 to 20. The multiplier and commercial layers were not explicitly simulated but were instead represented with the average genetic merit of nucleus males that would give rise to multiplication and commercial animals while accounting for the time lag. Specifically, we only focused on a breeding program that produced terminal males in a scheme that closely resembled a pig breeding program.

Simulations involved the following four steps:

(i) Generating genome,
(ii) Generating a quantitative trait and breeding values,
(iii) Generating an animal breeding program,
(iv) Selection and dissemination to the commercial layer with the conventional or surrogate sires strategy.

Results are presented as the mean of one hundred replicates for each scenario and encompass the genetic merit of nucleus males that would give rise to commercial animals at a given time point.

### Genome

Whole-genome sequences were generated using the Markovian Coalescent Simulator (MaCS) [12] and AlphaSim [13] for 400 base haplotypes for each of ten 10 chromosomes. Chromosomes (each 100 cM long and comprising 10^8^ base pairs) were simulated using a per site mutation rate of 2.5×10^−8^, a per site recombination rate of 1.0×10^−8^, and an effective population size (N_e_) that varied over time in accordance with estimates that are representative of livestock populations [e.g., 14–17] as follows: N_e_ was set to 100 in the final generation of the coalescent simulation, to N_e_ = 1256, 1000 years ago, to N_e_=4350, 10,000 years ago, and to N_e_=43,500, 100,000 years ago, with linear changes in between these time-points. The resulting sequences had approximately 540,000 segregating sites.

### Quantitative trait

A quantitative trait was simulated by randomly sampling 10,000 causal loci from the genome in the base population, with the restriction that 1,000 were sampled from each of the 10 chromosomes. For these loci, the allele substitution effect was randomly sampled from a normal distribution with a mean of 0 and standard deviation of 0.01 (1.0 divided by the square root of the number of loci).

### Breeding values

True breeding values were computed as a sum of effects at causal loci. To simulate selection without the full computational burden and complexity of simulating training sets and estimation with best linear unbiased prediction, we simulated pseudo estimates of breeding values by adding a level of noise to true breeding values. Different levels of noise were added to achieve a targeted accuracy. For the genomic tests we simulated accuracies of 0.50, 0.70 and 0.90. For the progeny tests we simulated accuracies as a function of the number of progeny [24] used in the different scenarios (described below).

### Breeding program

A pedigree of 42 generations for the nucleus population was simulated. Each generation included 1,000 (**SmallScenario**) or 5,000 (**BigScenario**) individuals with equal sex ratio. The different numbers of individuals were used to quantify impact of nucleus population size on the benefit of surrogate sire technology. All females (500 for the SmallScenario or 2,500 for the BigScenario) and 50 males were selected as the parents of each generation. This selection was based on a genomic test. In the first generation of the recent historical animal breeding population (i.e., generation −22), the chromosomes of each individual were sampled from the 400 base haplotypes. In later generations (i.e., generations −21 to 20), the chromosomes of each individual were sampled from parental chromosomes with recombination (assuming no interference). A recombination rate of 1 Morgan per chromosome was used, resulting in a 10 Morgan genome.

### Scenarios

Two different strategies were used to identify males from the nucleus who would give rise to commercial animals, either through conventional multiplication or surrogate sires. The conventional multiplication strategy used the top 50, 200, or 500 males in both the SmallScenario and the BigScenario. Males were chosen based on a genomic test. The surrogate sires strategy used multi-stage testing. Males were chosen based on an initial genomic test (S0), followed by one or two subsequent progeny tests (S1 and S2). As is the case with plant breeding programs, as the testing progressed through the stages we reduced the number of tested individuals and increased accuracy of tests. Based on the tests the best individual or set of individuals were identified and used as elite donors of spermatogonial stem cells to surrogate sires.

To quantify the impact of different amounts of testing resources and different allocation of these resources we simulated different accuracies of the genomic test at S0, different numbers of donor candidates tested with different number of progeny at S1 and S2. At S0 we simulated a genomic test with an accuracy of 0.50, 0.70, and 0.90. To ensure that each breeding program had the same costs, we assumed that a total of 14,000 progeny were available for progeny testing stages.

With single progeny test (S1) we used the 14,000 progeny to test 14 donor candidates each with 1,000 progeny, 28 donor candidates each with 500 progeny, 56 donor candidates each with 250 progeny, 112 donor candidates each with 125 progeny, 224 donor candidates each with 63 progeny, or 448 donor candidates each with 31 progeny.

With two progeny tests (S1 and S2) we used either 2,000, 4,000, or 6,000 progeny for the first test (S1) and the remaining 12,000, 10,000, or 8,000 for the second test (S2). At S1 either 100, 200, or 400 donor candidates were tested. Thus, when 2,000 progeny were used at S1 the 100, 200, or 400 donor candidates were each tested with 20, 10, or 5 progeny respectively. When 4,000 progeny were used at S1 the 100, 200, or 400 donor candidates were each tested with 40, 20, or 10 progeny respectively. When 6,000 progeny were used at S1 the 100, 200, or 400 donor candidates were each tested with 60, 30, or 15 progeny respectively. At S2 we tested either 10 or 20 donor candidates advanced from S1. When 12,000 progeny remained to be used at S2 the 10 or 20 donor candidates were each tested with 1,200 or 600 progeny respectively. When 10,000 progeny remained to be used at S2 the 10 or 20 donor candidates were each tested with 1,000 or 500 progeny respectively. When 8,000 progeny remained to be used at S2 the 10 or 20 donor candidates were each tested with 800 or 400 progeny respectively. From each of these testing strategies we chose either 1 or 5 donors of spermatogonial stem cells for surrogate sires in the commercial layer.

All of these different factors (two sizes of a breeding program [Small, Big], three conventional strategy scenarios [50, 200, 500 males], six surrogate sires strategy scenarios with two-stage testing, 18 surrogate sires strategy scenarios with three-stage testing, and using one or five donors) gave 102 different scenarios for each level of genomic test accuracy. The map of all these scenarios and used resources is summarized in Fig. 4.

### Time assumptions

The time taken to transfer germplasm from the nucleus to the commercial layer was assumed to be 3.5 years for the conventional strategy (but see the note below about “dilution”), 3.5 years for the surrogate sires strategy with two-stage testing, and 4.5 years for the surrogate sires strategy with three-stage testing. The different steps that underlie these time frames are presented in Fig. 3. We based our parameters on pigs and assumed 6 months for a male to reach sexual maturity, 4 months for a successful gestation, and 8 months to collect terminal line phenotypes on progeny. Based on these parameters we assumed 12 months to progeny test a sexually mature male. When the donors are identified we assumed that it takes a further 12 months to produce surrogate sires from these. Finally, we assumed a 12 months for the commercial progeny to pass through gestation and complete their growth. We assumed that the conventional program involved two rounds of multiplication that each take 12 months to complete.

Although we assumed that the genetic improvement with the conventional strategy is delivered to the commercial population in 3.5 years, we assumed an additional component of genetic lag, because the genetic merit of the sires entering the multiplier layer is “diluted” by the lagged genetic merit of females in the multiplier layer (i.e., we assumed no selection of females in the multiplier). Such a dilution would not occur with the surrogate sires strategy, because the multiplication layer does not arise. To account for this extra genetic lag in the conventional strategy we “diluted” genetic merit of commercial sires as follows:

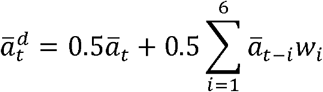

where *a̅_t_* is the average genetic merit of used nucleus males in generation *t* and *w_i_* is the relationship coefficient between the commercial sire and his maternal male ancestor in the generation *i*. We only accounted for 6 generations with *w_i_* ranging from 0.5 in *t* – 1 generation to 0.015625 in the *t* – 6 generation. This “dilution” increased genetic lag of the conventional strategy by an equivalent of ~1.04 years’ worth of extra genetic gain.

### Comparison of different scenarios

To ensure that sufficient numbers of generations had been traversed for “dilution”, we chose to present the results in terms of the genetic merit of terminal sires used in the commercial layer emerging from generation 11 and each subsequent generation. We report genetic merit in units of the standard deviation of true breeding values of the nucleus animals in the base generation (*σ_b_*), i.e., as (*a̅_t_* – *a̅_b_*) / *σ_b_*, where *a̅_t_* is the average true breeding value of the nucleus males that gave rise to commercial sires in year c and *a̅_b_* is the average true breeding value of nucleus animals in the base generation. Calculating the genetic merit of commercial sires in this way allowed the different strategies to be compared in terms of genetic merit of the commercial sires at the same year. Finally, we have converted the standardized genetic merit into years’ worth of genetic gain by calculating the number of years it takes the conventional breeding program when selecting the top 50 males to deliver the same level of genetic merit to the commercial layer.

## Results

The surrogate sires strategy increased the genetic merit of terminal sires used in the commercial layer. The genetic merit of commercial surrogate sires from the surrogate sires strategy was as much as 6.5 to 9.2 years’ worth of genetic gain higher than the genetic merit of commercial sires from the conventional multiplication strategy. In both the SmallScenario and BigScenario the three-stage testing strategy was the best strategy for identifying elite donors. The best performing three-stage testing strategy involved a genomic test at the first stage, 100 donor candidates tested each with 60 progeny at the second stage, and 20 donor candidates tested each with 400 progeny at the third stage (see Table 1 for details). The benefit of surrogate sires strategy was greatest when the genomic test accuracy was lowest and when the conventional strategy required large proportions of the nucleus males to be used for multiplication.

**Table 1.** Average Years’ worth of Genetic Gain (YGG) of the best performing surrogate sire strategy scenario above the conventional strategy that uses either 50, 200, or 500 males

| Genomic test accuracy | Males progeny tested S1 | Males progeny tested S2 | Progeny test resources<sup>1</sup> | Donors used | YGG<sub>50</sub> | YGG<sub>200</sub> | YGG<sub>500</sub> |
| --- | --- | --- | --- | --- | --- | --- | --- |
| Small Scenario |  |  |  |  |  |  |  |
| 0.5 | 100 | 20 | 6000S1 / 8000S2 | 1 | 6.5 | 7.5 | 9.2 |
| 0.7 | 200 | 20 | 6000S1 / 8000S2 | 1 | 4.5 | 6.5 | 7.2 |
| 0.9 | 200 | 20 | 6000S1 / 8000S2 | 1 | 2.4 | 4.5 | 5.0 |
| Big Scenario |  |  |  |  |  |  |  |
| 0.5 | 100 | 20 | 6000S1 / 8000S2 | 1 | 2.7 | 3.5 | 4.1 |
| 0.7 | 200 | 20 | 6000S1 / 8000S2 | 1 | 2.1 | 2.5 | 3.5 |
| 0.9 | 200 | 20 | 6000S1 / 8000S2 | 1 | 1.2 | 1.7 | 2.5 |

<sup>1</sup>Total number of progeny allocated in the first progeny test (S1) and in the second progeny test (S2)

In what follows the results are divided into three sub-sections for ease of presentation: (i) comparison of the conventional strategy and the best performing surrogate sires strategies; (ii) comparison of two-stage testing scenarios of the surrogate sires strategy; and (iii) comparison of three-stage testing scenarios of the surrogate sires strategy. To avoid clutter in the figures or tables we do not show standard errors across the 100 replicates of the simulated scenarios because the standard errors were small in all instances less than 0.009 YGG.

### Comparison of the conventional and the best performing surrogate sires strategies

Fig. 5 and S1 show the average genetic merit of commercial sires derived from the best performing surrogate sires strategy scenario and the conventional strategy against time, for three different genomic test accuracies (0.5, 0.7, and 0.9) and the SmallScenario and the BigScenario. The conventional strategy used the top 50, 200, or 500 males in multiplication. At all points in time and for all three genomic accuracies commercial sires derived from the best performing surrogate sires strategy scenario had a higher genetic merit than those derived from the conventional strategy. This benefit was greater when more males were used for multiplication in the conventional strategy. The benefit of using surrogate sires strategy decreased as the genomic test accuracy increased. Across time the difference between the two strategies was almost constant. These trends were common both in the SmallScenario and the BigScenario, although with differences in magnitude.

**Fig. 5.**
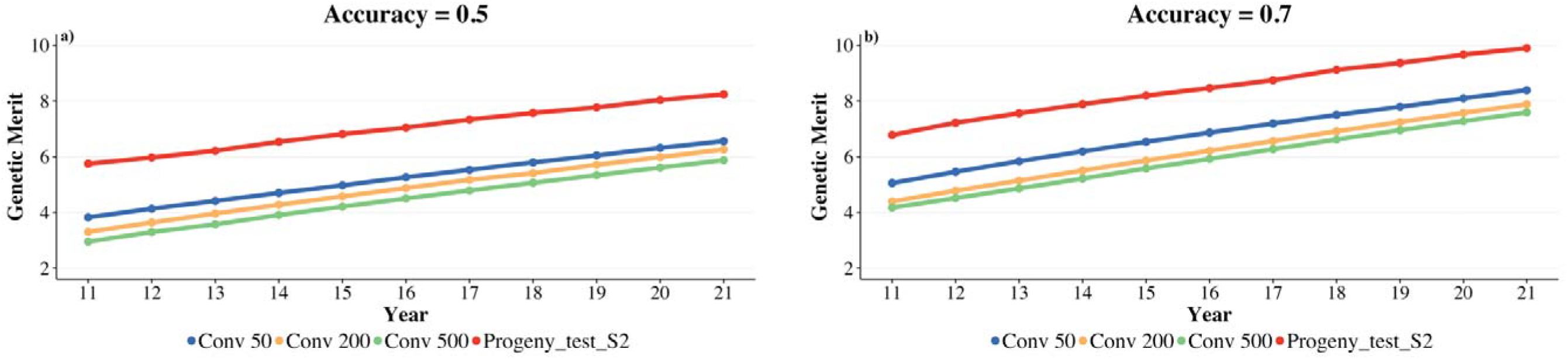
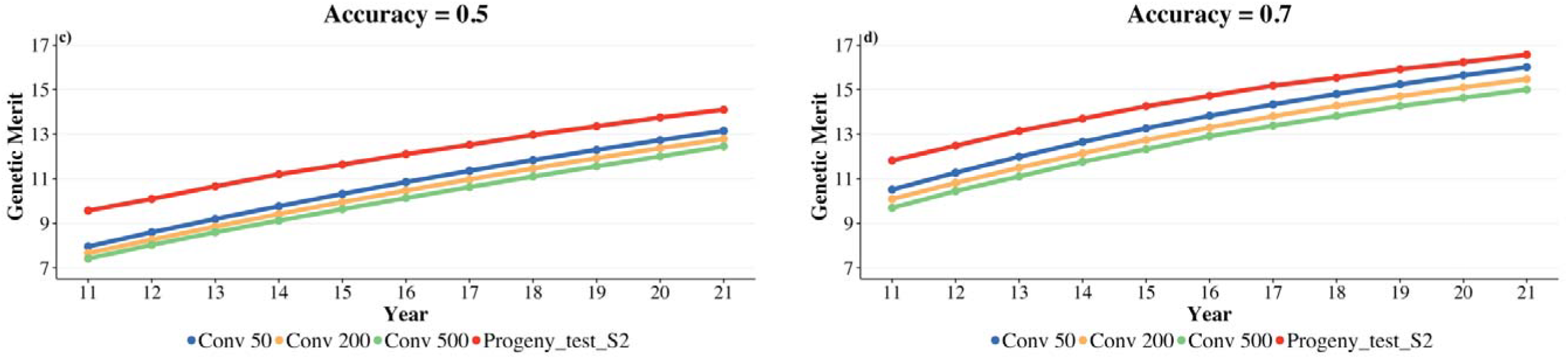
Average genetic merit of commercial sires derived from the best performing surrogate sire strategy scenario and the conventional strategy (top 50, 200 and 500 males) for SmallScenario (a and b) and BigScenario (c and d) plotted against time

Table 1 enumerates some of the main results than can be observed in Fig. 4 and S1. Across all scenarios tested the best performing surrogate sires strategy scenario involved first a genomic test of all donor candidates followed by two subsequent progeny tests and the use of a single elite donor. The benefit of surrogate sires strategy above the conventional strategy was greater when more males were used for multiplication with the conventional strategy. When the genomic test accuracy was low (0.5) the best strategy was to first progeny test 100 candidates on 6,000 progeny and then to test 20 candidates on 8,000 progeny. This testing and subsequent production of surrogate sires was assumed to take one additional year compared to the conventional strategy. After accounting for this extra time and for the dilution in the conventional multiplication process, we observed that in the SmallScenario the surrogate sires strategy delivered on average between 6.5 and 9.2 years’ worth of extra genetic gain in commercial sires compared to the conventional strategy that uses respectively between 50 and 500 males in multiplication. For the BigScenario the equivalent values were between 2.7 and 4.1 years’ worth of extra genetic gain.

When the genomic test accuracy was higher (> 0.5) the optimal allocation of testing resources was slightly different. Instead of first progeny testing 100 candidates, as was the case when the genomic test accuracy was low, progeny testing 200 candidates was the best performing scenario. All other scenario parameters were the same as when the genomic test accuracy was low. The benefit of surrogate sires strategy decreased with the increasing genomic test accuracy and the magnitude of benefit differed significantly between the SmallScenario and the BigScenario (Table 1).

On average the surrogate sires strategy in SmallScenario delivered between 6.5 and 9.2 years’ worth of extra genetic gain in commercial sires when the genomic test accuracy was 0.5. When the genomic test accuracy was 0.7 these values reduced to between 4.5 and 7.2 years and when the genomic test accuracy was 0.9 they further reduced to between 2.4 and 5.0 years.

On average the surrogate sires strategy in BigScenario delivered between 2.7 and 4.1 years’ worth of extra genetic gain in commercial sires when the genomic test accuracy was 0.5. When the genomic test accuracy was 0.7 these values reduced to between 2.1 and 3.5 years and when the genomic test accuracy was 0.9 they further reduced to between 1.20 and 2.50 years.

The differences in the SmallScenario and the BigScenario were due to the different proportions of males used in multiplication to give rise to commercial sires. In the SmallScenario 10% to 100% of males were used while the in the BigScenario 2% to 20% of males were used.

For simplicity of presentation and based on the consistency of trends described above, in the following sections we only present comparisons to the conventional strategy in which 50 males were used in multiplication.

### Comparison of two-stage testing scenarios of the surrogate sires strategy

Tables 2 and 3 show the performance of different two-stage testing scenarios of the surrogate sires strategy. Performance is measured as the average years’ worth of extra genetic gain in the commercial sires delivered by the surrogate sires strategy compared to the conventional strategy for both the SmallScenario (Table 2) and the BigScenario (Table 3). Consistent with the results reported in the previous sub-section the benefit of surrogate sires strategy was always lower when the genomic test accuracy was higher. In some scenarios, the benefit was minimal. In all cases, there was an intermediate optimum for the numbers of candidates tested. Using five elite donors was always worse than using one. This behaviour was observed in both the SmallScenario and the BigScenario although with some interesting differences. The BigScenario showed a general shrinkage of years’ worth of genetic gain compared to the SmallScenario, resulting in a general increase in the number of scenarios that showed a small benefit of the surrogate sires strategy.

**Table 2.** Average Years’ worth of Genetic Gain (YGG) with the two-stage testing scenarios of the surrogate sire strategy above the conventional strategy that uses 50 males (SmallScenario)

| Males Tested | Progeny/Male | Donors used | YGG<sub>0.5</sub><sup>1</sup> | YGG<sub>0.7</sub><sup>1</sup> | YGG<sub>0.9</sub><sup>1</sup> |
| --- | --- | --- | --- | --- | --- |
| 14 | 1000 | 1 | 4.1 | 3.0 | 1.8 |
| 28 | 500 | 1 | 4.7 | 3.0 | 1.2 |
| 56 | 250 | 1 | 5.1 | 3.5 | 2.2 |
| 112 | 125 | 1 | 5.3 | 3.6 | 2.2 |
| 224 | 63 | 1 | 4.8 | 2.8 | 1.3 |
| 448 | 31 | 1 | 3.8 | 2.1 | 1.1 |
| 14 | 1000 | 5 | 2.9 | 1.9 | 0.2 |
| 28 | 500 | 5 | 3.1 | 2.1 | 0.5 |
| 56 | 250 | 5 | 3.6 | 2.4 | 1.1 |
| 112 | 125 | 5 | 3.6 | 2.6 | 1.2 |
| 224 | 63 | 5 | 3.4 | 1.9 | 0.3 |
| 448 | 31 | 5 | 2.8 | 1.6 | 0.2 |

<sup>1</sup>Genomic test accuracy at the initial stage (S0) 0.5, 0.7, and 0.9

**Table 3.** Average Years’ worth of Genetic Gain (YGG) with the two-stage testing scenarios of the surrogate sire strategy above the conventional strategy that uses 50 males (BigScenario)

| Males Tested | Progeny/Male | Donors used | YGG<sub>0.5</sub><sup>1</sup> | YGG<sub>0.7</sub><sup>1</sup> | YGG<sub>0.9</sub><sup>1</sup> |
| --- | --- | --- | --- | --- | --- |
| 14 | 1000 | 1 | 2.3 | 1.7 | 0.7 |
| 28 | 500 | 1 | 2.4 | 1.9 | 0.8 |
| 56 | 250 | 1 | 2.5 | 2.0 | 1.0 |
| 112 | 125 | 1 | 2.5 | 2.0 | 1.1 |
| 224 | 63 | 1 | 2.0 | 1.8 | 0.8 |
| 448 | 31 | 1 | 1.9 | 1.5 | 0.4 |
| 14 | 1000 | 5 | 1.7 | 1.2 | 0.5 |
| 28 | 500 | 5 | 1.7 | 1.2 | 0.7 |
| 56 | 250 | 5 | 1.9 | 1.1 | 1.0 |
| 112 | 125 | 5 | 2.0 | 1.1 | 1.0 |
| 224 | 63 | 5 | 1.8 | 1.0 | 0.5 |
| 448 | 31 | 5 | 1.0 | 0.8 | 0.3 |

<sup>1</sup>Genomic test accuracy at the initial stage (S0) 0.5, 0.7, and 0.9

At all levels of genomic test accuracy the best scenario was to screen candidates based on genomic test, progeny test 112 candidates each with 125 progeny, and use the best candidate as a single elite donor. With the genomic test accuracy of 0.5, 0.7, and 0.9 this scenario gave respectively 5.3, 3.6, or 2.2 years’ worth of extra genetic gain in commercial sires in the SmallScenario (Table 2) and respectively 2.5, 2.0 or 1.1 years’ worth of extra genetic gain in commercial sires in the BigScenario (Table 3).

Just as for the case of selecting one elite donor of spermatogonial cells for surrogate sires, when selecting five elite donors, progeny testing 112 candidates each with 125 progeny gave the highest benefit. With the genomic test accuracy of 0.5, 0.7, and 0.9 this scenario gave respectively 3.6, 2.6 and 1.2 years’ worth of extra genetic gain in the SmallScenario (Table 2) and respectively 2.0, 1.1 and 1.0 in the BigScenario (Table 2).

### Comparison of three-stage testing scenarios of the surrogate sires strategy

Tables 4 and 5 shows the performance of different three-stage testing scenarios of the surrogate sires strategy in the SmallScenario when either one or five elite donors used. By varying several parameters, we tested 216 (108 for the SmallScenario and 108 for the BigScenario) different scenarios of three-stage testing with fixed total progeny testing resources. These resources were the same as for the two-stage testing scenarios described in the previous sub-section. The parameters with the three-stage testing scenarios were the genomic test accuracy for the first stage, the split of resources between the two subsequent progeny tests, the number of tested donor candidates, the number of progeny per tested donor candidate at each progeny test stage, and the number of elite donors used for production of surrogate sires.

**Table 4.** Average Years’ worth of Genetic Gain (YGG) with the three-stage testing scenarios of the surrogate sire strategy with one elite donor above the conventional strategy that uses 50 males (SmallScenario)

| Progeny test resources<sup>1</sup> | Males progeny tested S1 | Progeny/Male S1 | Males progeny tested S2 | Progeny/Male S2 | YGG<sub>0.5</sub><sup>2</sup> | YGG<sub>0.7</sub><sup>2</sup> | YGG<sub>0.9</sub><sup>2</sup> |
| --- | --- | --- | --- | --- | --- | --- | --- |
| 2000S1/12000S2 | 100 | 20 | 10 | 1200 | 5.3 | 3.5 | 2.2 |
|  |  |  | 20 | 600 | 5.4 | 3.6 | 2.4 |
|  | 200 | 10 | 10 | 1200 | 4.9 | 3.2 | 2.2 |
|  |  |  | 20 | 600 | 5.1 | 3.3 | 2.1 |
|  | 400 | 5 | 10 | 1200 | 4.5 | 3.7 | 2.0 |
|  |  |  | 20 | 600 | 4.7 | 2.7 | 1.4 |
| 4000S1/10000S2 | 100 | 40 | 10 | 1000 | 5.5 | 3.6 | 2.2 |
|  |  |  | 20 | 500 | 5.8 | 4.0 | 2.3 |
|  | 200 | 20 | 10 | 1000 | 5.3 | 3.5 | 2.4 |
|  |  |  | 20 | 500 | 5.4 | 3.8 | 2.3 |
|  | 400 | 10 | 10 | 1000 | 4.3 | 3.3 | 1.6 |
|  |  |  | 20 | 500 | 4.5 | 3.5 | 1.4 |
| 6000S1/8000S2 | 100 | 60 | 10 | 800 | 5.9 | 4.1 | 2.0 |
|  |  |  | 20 | 400 | 6.5 | 4.2 | 2.2 |
|  | 200 | 30 | 10 | 800 | 5.3 | 4.2 | 2.1 |
|  |  |  | 20 | 400 | 5.7 | 4.5 | 2.4 |
|  | 400 | 15 | 10 | 800 | 5.0 | 3.4 | 1.6 |
|  |  |  | 20 | 400 | 5.8 | 3.5 | 1.2 |

<sup>1</sup>Number of total progeny allocated in the first progeny test (S1) and in the second progeny test(S2)
<sup>2</sup>Genomic test accuracy at the initial stage (S0) 0.5, 0.7, and 0.9

**Table 5.** Average Years’ worth of Genetic Gain (YGG) with the three-stage testing scenarios of the surrogate sire strategy with five elite donors above the conventional strategy that uses 50 males (SmallScenario)

| Progeny test resources<sup>1</sup> | Males progeny tested S1 | Progeny/Male S1 | Males progeny tested S2 | Progeny/Male S2 | YGG<sub>0.5</sub><sup>2</sup> | YGG<sub>0.7</sub><sup>2</sup> | YGG<sub>0.9</sub><sup>2</sup> |
| --- | --- | --- | --- | --- | --- | --- | --- |
| 2000S1/12000S2 | 100 | 20 | 10 | 1200 | 4.1 | 2.1 | 1.1 |
|  |  |  | 20 | 600 | 4.4 | 2.2 | 1.2 |
|  | 200 | 10 | 10 | 1200 | 3.0 | 2.1 | 1.2 |
|  |  |  | 20 | 600 | 3.7 | 2.5 | 1.3 |
|  | 400 | 5 | 10 | 1200 | 2.2 | 1.5 | 1.0 |
|  |  |  | 20 | 600 | 2.2 | 1.4 | 1.0 |
| 4000S1/10000S2 | 100 | 40 | 10 | 1000 | 4.4 | 2.4 | 1.3 |
|  |  |  | 20 | 500 | 4.5 | 2.5 | 1.2 |
|  | 200 | 20 | 10 | 1000 | 4.1 | 2.2 | 1.1 |
|  |  |  | 20 | 500 | 4.1 | 2.7 | 1.2 |
|  | 400 | 10 | 10 | 1000 | 4.2 | 1.7 | 1.0 |
|  |  |  | 20 | 500 | 4.2 | 2.0 | 1.8 |
| 6000S1/8000S2 | 100 | 60 | 10 | 800 | 4.5 | 3.1 | 1.6 |
|  |  |  | 20 | 400 | 5.0 | 3.2 | 1.8 |
|  | 200 | 30 | 10 | 800 | 4.6 | 2.1 | 1.3 |
|  |  |  | 20 | 400 | 5.0 | 2.2 | 1.4 |
|  | 400 | 15 | 10 | 800 | 4.1 | 1.7 | 1.2 |
|  |  |  | 20 | 400 | 4.6 | 2.2 | 1.2 |

<sup>1</sup>Number of total progeny allocated in the first progeny test(S1) and in the second progeny test(S2)
<sup>2</sup>Genomic test accuracy at the initial stage (S0) 0.5, 0.7, and 0.9

The three-stage testing gave a greater benefit than the two-stage testing. As for the two-stage testing, using one elite donor for surrogate sires gave a greater benefit than using five elite donors and the benefit of surrogate sires strategy was greater when the genomic test accuracy was lower. A total of 14,000 progeny were split across the two stages of progeny testing. Increasing the resources in the first progeny test increased benefit of surrogate sires strategy. For example, with the SmallScenario when the genomic test accuracy was 0.5, 6,000 progeny were used in the first progeny test, 8,000 were used in the second progeny test, and when one elite donor was used in the end, the benefit was 6.5 years’ worth of extra genetic gain in commercial sires above the conventional strategy that uses 50 nucleus males in multiplication. This was a greater benefit than the 5.8 years’ worth of extra genetic gain for the scenario that split the 14,000 progeny into 4,000 for the first progeny test and 10,000 for the second progeny test, which was in turn better than the 5.4 years’ worth of extra genetic gain for the scenario that split the 14,000 progeny into 2,000 for the first progeny test and 12,000 for the second progeny test. This trend of greater benefit when more progeny were dedicated to the first progeny test was observed for almost all tested scenarios.

For the SmallScenario the difference between testing 100 or 200 donor candidates at the first progeny test was not consistent. That said, when the genomic test accuracy was 0.5, allocating 100 candidates to the first progeny test was usually better than allocating 200, and allocating 200 candidates was usually better than allocating 400. At higher genomic test accuracies, there were little differences between allocating 100 or 200 candidates to the first progeny test, but both of these sets of scenarios were usually better than allocating 400 candidates to the first progeny test.

In the SmallScenario allocating 20 elite donor candidates to the second progeny test was almost always better than allocating 10 candidates. A total of 54 scenarios were evaluated for SmallScenario. In only 6 of these scenarios allocating 10 candidates was better than allocating 20.

Overall for the SmallScenario, when the genomic test accuracy was 0.5, the best three-stage testing scenario used 6,000 progenies in the first progeny test of 100 candidates each with 60 progeny, 8,000 progenies in the second progeny test of 20 candidates each with 400 progeny, and used a single elite donor for surrogate sires. This scenario gave a benefit of 6.50 years’ worth of extra genetic gain in commercial sires compared to the conventional strategy. The same distribution of testing resources was also the joint best when five, instead of one, elite donors were used for surrogate sires.

The same trends as for the SmallScenario were observed also for the BigScenario, but with smaller benefit of the surrogate sire strategy (See table S1 and S2).

## Discussion

The results of this paper suggest that a surrogate sires strategy could be very beneficial for the dissemination of genetic gain in animal breeding. In summary, our results indicate that benefits of the as much as 6.5 to 9.2 years’ worth of genetic gain in commercial sires could be realized with surrogate sires compared to the conventional multiplication. It was best to identify elite donors for surrogate sires via a three-stage testing strategy involving a first screen with a genomic test followed by two subsequent progeny tests. The benefits of a surrogate sires strategy were greater when genomic test accuracy was low and when the conventional strategy used a large proportion of males in multiplication. To discuss these results we divide the discussion into four sections: (i) possible explanations for the observed trends; (ii) justification and impact of assumptions; (iii) the potential impact of surrogate sires on the redesign of animal breeding programs; and (iv) risks and opportunities of using surrogate sires.

### Possible explanations for the observed trends

That surrogate sire technology generates such a benefit in terms of years’ worth of genetic gain can be explained in the context of the breeders’ equation. While the surrogate sires strategy does not rely on the selection of the best individuals and using them as parents of the next generation, it does rely on the identification of the best individuals from a cohort and using them as donors of spermatogonial cells for surrogate sires, which is another form of the selection problem. In any cohort, the best few individuals will be some number of standard deviations above the cohort average. For example, when surrogate sires technology delivered 6.5 years’ worth of additional genetic gain in commercial sires the best nucleus male was on average 2.7 standard deviations above the cohort mean. In contrast, the best 50 nucleus males were 2.0 standard deviations above the cohort mean. Given that the breeding program proceeded at a rate of genetic progress of 0.4 standard deviations per year, choosing the best male as a donor for surrogate sires rather than the best 50 males produced surrogate sires that were better for more than 8 years’ worth of genetic gain. However, accounting for the imperfect accuracy of identifying donors with the surrogate strategy or the best 50 males for multiplication with the conventional strategy and the time to generate commercial sires with either strategy the final result was 6.5 years’ worth of genetic gain.

With constant progeny test accuracies the benefit of the surrogate sires strategy depended on the proportion of male candidates that the conventional strategy used to give rise to commercial sires. If the breeding program needed to use a large proportion of its nucleus male candidates (e.g., the best 200 or 500) the benefit of surrogate sires strategy was greater than if it needed to use a few. Again, this result is entirely consistent with the breeders’ equation. Specifically, it can be explained in the context of selection intensity being a nonlinear function of the percentage of selected individuals, i.e., selection intensity increases almost linearly down to 20 or 10% selected, but increases sharply (nonlinearly) thereafter. While both conventional and surrogate sires strategies exploit the tail of distribution with high selection intensities, the surrogate sires strategy also exploits the steeper part. This explains why the benefit of surrogate sires was higher in the SmallScenario than in the BigScenario. In the SmallScenario we had 500 candidates and selected 100 with the conventional strategy (percentage selected 20% and selection intensity 1.4) or 1 with the surrogate sires strategy (percentage selected 0.2% and selection intensity 3.2). In the BigScenario we had 2,500 candidates and selected 100 with the conventional strategy (percentage selected 4% and selection intensity 2.2) or 1 with the surrogate sires strategy (percentage selected 0.04% and selection intensity 3.6). The same logic also explains why selecting five as opposed to one donor for surrogate sires gave a lower benefit.

The observed differences in the performance of different surrogate sire strategies can also be explained in the context of the breeders’ equation. When the genomic test accuracy used in the first stage of testing was lower the benefit of surrogate sires strategy was higher. Under the conventional strategy, the average genetic merit of the nucleus males that gave rise to commercial sires was lower when the genomic test accuracy was lower than when it was higher. With surrogate sires strategy this reduction in genetic merit due to the low genomic test accuracy is compensated by the subsequent progeny tests. This is in line with the analysis of Dickerson and Hazel [18], who compared the use of progeny test as a supplement to earlier culling on own or sibling performance. Their conclusion was that progeny testing is warranted when heritability is low in which case accuracy of estimated breeding values from own or sibling phenotypes (or genomic prediction in our study) is low. Genomic selection can be thought of as a light touch first screen, the purpose of which is to identify the top group of animals, which are then tested on many progeny. The purpose of subsequent progeny tests is then a search for the best individual within this group.

This same logic also explains why the three-stage testing was better than the two-stage testing. Both schemes started with a genomic test that was followed by one progeny test with the two-stage testing or two subsequent progeny tests with the three-stage testing. With the three-stage testing the first progeny test serves to use a portion of resources to evaluate many candidates relatively accurately in order to discard most candidates. Then the second progeny test uses the remaining resources to even more accurately identify the final candidate. In terms of the breeders’ equation the first progeny test maximizes selection intensity, while the second maximizes accuracy. The three-stage testing appears to address both of these parameters more optimally than the two-stage testing.

There is a substantial body of literature on multi-stage selection [19–23] which the observed trends in this study are consistent with. It is well known that increasing the number of progeny per candidate increases accuracy [24,25] and that the number of candidates to be tested is important and the trade-off between the two must be found. In our simulations, we found the optimum at progeny testing 112 candidates, given a fixed amount of resources, in our case 14,000 progeny. This optimum was consistent across the different levels of genomic test accuracy. However, the level of genomic test accuracy heavily influenced the amount of extra genetic gain, because higher accuracy directly translates to higher genetic gain. These trends are consistent with the long-established multi-stage testing in plant breeding [9]. Most plant breeding programs use multi-stage testing to identify elite single genotype (e.g., inbred line) that is then deliver to the commercial layer. Typically, these programs initially screen many individuals imprecisely at the first stage. At each subsequent stage they reduce the number of tested individuals, but the testing precision is increased.

### Justification and impact of assumptions

There is a huge range of possible strategies for the identification of donors for surrogate sires and we only evaluated a small subset in this study. We choose the tested range of scenarios because we believe they could demonstrate the properties of surrogate sires strategy. They show that in some circumstances surrogate sires can deliver a large benefit and in others small benefit. We chose the three levels of genomic test accuracy as these levels reflect what might be possible in breeding programs of various sizes. To ensure that all strategies used an equal set of resources we set the total number of progeny involved in progeny testing to 14,000. We chose this number as it was divisible in many ways and thereby enabled several strategies to be compared and because this number was similar the 10,000 progeny that would be used by an animal breeding program that each year tested 100 candidates each with 100 progeny, a scale of progeny testing that was not uncommon in some animal breeding programs before the advent of genomic selection.

With the two-stage testing the total testing resources were distributed across many or few candidates. As expected, testing an intermediate to high number of candidates (i.e., 112 to 224) on a relatively small number of progeny (i.e., 125 to 63) gave higher benefits than testing a few candidates (e.g., 14) on many progeny or a very high number of candidates (448) on few progeny (31). These trends fit the expectations from the breeders’ equation and occur due to the interplay between selection intensity and accuracy. However, when the chosen elite donors of spermatogonial cells for surrogate sires are to be used to produce huge numbers of progeny in the commercial layer, the risk of a donor carrying some major defect that was not identified by the testing process must also be minimized. For this reason, it is unlikely that a strategy in which donors are tested with a single stage of progeny testing using a ~200 or less progeny would ever be used by a commercial breeding program.

It was this logic that motivated us in our design of the three-stage testing scheme. Our intuition was that the first progeny test would evaluate many candidates with relatively low accuracy, while the second progeny test would evaluate a handful of individuals with high accuracy, i.e., 10 or 20 candidates each with respectively 800 or 400 progeny. Using many progeny ensures high accuracy, but also a high degree of certainty that the final donor(s) would not carry any major defects.

A major assumption of this study was the amount of time it took to identify elite donors and then to make surrogate sires. It is likely that the different time assumptions could be shortened or lengthened for both the conventional multiplication strategy and the surrogate sires strategy in several ways and depending on the assumed species. The benefit of surrogate sires strategy would change accordingly.

Finally, we choose to model a pig breeding program in this study because this is the species that we are most familiar with. The benefits may be greater or smaller for other species. The benefits depend on the ratio of existing reproductive rates of males versus that enabled by surrogate sire technology, the time and cost associated with performing progeny tests, the levels of accuracy that can be obtained by genomic prediction and the relative cost and technical possibilities of surrogate sire technology itself in a particular species. Incidentally, in this study we did not account for the cost aspects of surrogate sire technology itself. Undoubtedly developing the technology itself will be hugely expensive and these costs of development may impact its eventual commercial cost. That said, in time many biotechnologies which are initially expensive become much cheaper (e.g., nowadays genotyping and animal cloning are both relatively inexpensive compared to their former costs) and we anticipate that surrogate sire technology will follow a similar pattern. However, given we have ignored the cost component of surrogate sire technology its benefit may be overestimated based on our results compared to a study which would account for such costs.

### The potential impact of surrogate sires on the redesign of animal breeding programs

Animal breeding programs maximize the genetic merit of commercial animals within the available financial, physical, technical, and physiological constraints. Implicitly a breeding program has two objectives: (i) improving the mean of the population; and (ii) delivering a product to the commercial producers. In dairy cattle for example, before the advent of genomic selection, breeders used progeny testing schemes that intensively evaluated relatively small numbers of candidate males and used the best of these as parents to improve the population, but also as a commercial and breeding product to be used by the commercial layer. In doing so, dairy cattle breeders maximised selection accuracy, but were constrained in their ability to increase selection intensity and decrease generation interval. However, commercial producers used well tested sires and therefore an individual producer could rely on using relatively few sires, who together could serve entire geographic regions. The advent of genomic selection changed this paradigm. Under genomic selection progeny testing of a small number of candidates has been replaced with a genomic testing of a large number of candidates. Those with best predictions are used as parents to improve the “open” nucleus population, but are also sold to commercial layer as a team of sires product (i.e., a group of sires sold together rather than a single sire sold on its own). In doing so, dairy cattle breeders increased selection intensity and reduced generation interval, but are constrained in their ability to achieve very high accuracy. Given that each candidate male has not had their merit assessed based on phenotypes of their progeny, there is a risk that certain sires are not that good or may carry mutations that are highly detrimental (e.g., a *de-novo* mutation that prevents progeny from lactating) [26–28]. To overcome this risk, breeders recommend that commercial producers use semen of a larger number (i.e., a team) of sires and limit their use of any one sire.

A surrogate sires strategy would need to exploit aspects of both genomic and progeny testing. Genomic testing can be used to drive the population improvement and, as demonstrated in the present paper, to identify a set of candidates that could enter a progeny testing scheme as part of the product development. The role of the progeny testing is to ensure that the chosen elite donors that give rise to surrogate sires released to the commercial sector are good animals, that they are not significantly worse than it is predicted by a genomic test and that they do not carry detrimental mutations. As demonstrated by the results of the present study two subsequent progeny tests used resources more efficiently than a single progeny test. Such multi-stage testing has a long history of use in plant breeding which also has a long history delivering products to commercial producers in a way that is highly analogous to what surrogate sires would enable for animal producers.

The majority of commercial producers for all of the major crops (maize, wheat, rice) use inbred lines or their hybrids. These inbred or hybrid lines can be grown on huge areas. Plant and animal breeding designs have diverged somewhat over the years owing to differences in biology, economics, and technical possibilities. Surrogate sire technology, combined with genomic selection, could result in a coalescence of designs across plant and animal breeding. One such design that could apply to both is the two-part scheme recently proposed by Gaynor et al. (2017) [11]. In this scheme, rapid recurrent selection based on genomic testing is used to increase the mean of the population, while multi-stage testing (genomic and phenotypic) is used to periodically extract, test, and develop a product from the population. The population improvement component resembles the nucleus of animal breeding programs, while the product development component resembles the multi-stage testing to derive inbred or hybrid lines of plant breeding programs. The latter could also be seen as an improved multiplication layer of animal breeding programs that exploit breed complementarity to deliver a commercial product.

In the present work, we focused on the use of surrogate sires to produce commercial animals (e.g., a terminal sire in a pig population). To do this, donors for surrogate sires were chosen based on their general combining ability. The strategy could also be extended to exploit specific combining ability to produce a relatively homogenous set of females from a maternal line that are crossed with single terminal male (via surrogate sires). Use of specific combining ability is widespread in hybrid crops where it exploits complementarity of pairs of individuals and heterosis generated by specific pairs of individuals. The surrogate sires strategy proposed in the present paper could be extended to exploit specific combining ability by adding additional stages that progeny tests specific crosses as is conducted in hybrid plant breeding programs. Because in livestock the parents are outbred (compared to crops where they are often inbred), a tiered strategy may be needed in the maternal line(s) that homogenizes dam haplotypes. For example, using a single surrogate sire, grandsire, and great-grandsire on the maternal population would give a pool of females that carried one of two haplotypes for 87.5% (0.5 + 0.25 + 0.125) of their genome. The terminal surrogate sire would be chosen based on a specific combining ability to these haplotypes.

### Risks and opportunities of using surrogate sires

Surrogate sires present risks and opportunities to commercial production. The most obvious risk relates to the genetic homogeneity of commercial animals if a single surrogate sire, or a set of very closely related surrogate sires were used. If a disease emerged that this homogenous group of animals was susceptible to, it could have a major detrimental impact on the commercial production. Having such large groups of homogeneous animals would also increase the selection pressure on disease pathogens to evolve pathogenicity to the group. Plant breeders and commercial crop growers have extensive experience in managing the potential to have genetic homogeneity across large segments of the production area. They have developed strategies to minimize the risk of disease outbreaks and other failures such as crop rotation, using multiple varieties on a farm, creating varietal blends consisting of multiple genotypes, and taking holistic strategies to pathogen management [29]. Aside from rotation, which is practically impossible in the animal sector, these strategies might have important roles in ensuring the effectiveness of surrogate sires in livestock.

A further risk of the homogenisation of the commercial population relates to genetic diversity. The genetic diversity contained in current populations is potentially a useful reserve of genetic diversity that could be used in breeding programs in case the nucleus genetic diversity was to become inappropriate at some point in the future (e.g., due to a disease catastrophe or because it became exhausted). Homogenisation of the commercial population would remove this safety net requiring greater care to be taken in the preservation of genetic diversity. Genebanks using frozen semen, eggs, or embryos are well established ways to preserve genetic diversity. There are also new ways which include the use of cultured primordial germ cells [30].

Undetected but highly deleterious mutations also pose a risk for the use of surrogate sires. While it is unlikely that this would arise after sufficient testing, it is not impossible. One such route could be through the occurrence of one or more such mutations arising as somatic mutations after the animal had been tested, leading to a mosaicism, which might affect sets of surrogate sires from the donor.

The most obvious opportunity emanating from surrogate sires also relates to the genetic homogeneity of commercial animals and can also draw on practices that are well established in crop production. In crops, management plans are supplied to a farmer alongside the seed (e.g., https://catalog.extension.oregonstate.edu/em9004). These plans are specifically tailored to the variety genotype based on extensive sets of field trials. They include recommendations for target market, expected performance, optimum sowing date, seeding rate, soil type and water, fertilizer, pesticide and fungicide requirements. These management plans complement the genetics of the variety and increase the benefit obtained from the genetic potential in a generic environment. Similar management plans could be developed for surrogate sires and the benefits would be similarly expected to exceed the benefit that was observed in the present study for the genetics alone (e.g., 6.5 year’s worth of genetic gain). The phenotype data collected to development of the management plans would also serve to further test and validate a particular donor.

Another obvious opportunity emanating from surrogate sires that also relates to the genetic homogeneity is the potential for increasing the product homogeneity. In animal production, product uniformity is an important topic. In meat animals, for instance, uniformity has economic benefits because excessive variability in carcass weight or conformation is penalized by slaughterhouses [31,32]. A genetically homogeneous commercial population, achieved through the use of surrogate sires, could aid product uniformity. However, if this was to be achieved, most of the increase in uniformity would need to emanate from matching very specific management plans to the homogenous genetics because homogeneous genetics in itself has limited ability to increase phenotypic homogeneity. Van Vleck [33] showed that in the context of cloned animals, if heritability is 25%, then the phenotypic standard deviation among clones would be 87% of that of uncloned animals and only if heritability is 100%, will clone mates have complete uniformity.

Compared to the conventional multiplication the surrogate sires strategy enables shorter lag between nucleus and commercial layer and requires a smaller number of parents contributing to the commercial layer. This offers several advantages including the ability to rapidly change the entire genetics in the commercial layer. This could be used to rapidly respond to sudden changes in requirements such as pressure from a new disease or the emergence of a new market for the product that has specific requirements (e.g., meat marbling).

The surrogate sire strategy would be costly to implement in practice because it would require capacity in advanced molecular biology and infrastructure for progeny testing. However, it presents other opportunities through which costs can be saved. For example, multiplier populations to produce terminal sires would not need to be maintained. This would free up resources for other investment in breeding programs, such as more progeny testing of donor candidates.

The surrogate sires strategy presents breeding programs with an enhanced opportunity to protect its intellectual property via limited release of males (thereby limiting the access of competitors to the broader source germplasm) and by exploitation of specific combining ability. This protection would give the breeding companies incentive to invest more and help to avoid the commonly observed market failure in some breeding industries. When intellectual property is properly protected, breeding companies are anecdotally reported to share the benefits two-thirds to the farmers and one third to the breeding company. Such sharing more than offsets the purchase cost to a producer, while it also gives profit to the breeder. Perhaps the most spectacular example of the benefits of such ways to reward investment in intellectual property are seen in maize which has seen a 6-fold increase in productivity since hybrid breeding was introduced in the 1930’s [34]. By releasing hybrids breeding organisations can protect the intellectual property that is their source germplasm. This in turn enables them to invest heavily in breeding activities (e.g., technology, field testing networks) that in turn drive accelerated genetic gains.

At least two barriers exist that may prevent the deployment of this technology in in real livestock breeding program. Firstly, genome editing currently appears to be the technology that is most likely to enable genome editing to be implemented in practice [2]. Globally, the future of governmental regulation of genome editing technology is currently uncertain which places uncertainty on the possibility for practical implementation of surrogate sire technology in real livestock breeding program. Secondly, effective deployment of surrogate sire technology will require partitioning of animal breeding programs into population improvement and product development parts. Product development will require deployment of extensive progeny testing schemes. Over the past decade the advent of genomic selection has removed progeny testing schemes from many breeding programs. Reinstating such schemes would be costly and further work will be needed to demonstrate the exact return on investment.

Finally, the results of this study raise an important question for existing breeding programs that use artificial insemination for dissemination. As noted above, genomic selection has led to the removal of progeny testing schemes from many livestock breeding programs. Our results raise some doubts about the merit of this. They show that when a breeding program releases a small number of individuals that are deployed widely there is a benefit to progeny testing these individuals. The degree of benefit depends on the accuracy of genomic selection, the number of individuals released and their subsequent usage, and the accuracy and the number of stages in a progeny testing scheme and the relative time taken to perform a progeny test. Determining whether the removal of progeny testing schemes from genomic selection driven livestock breeding programs was the right thing to do in retrospect is beyond the scope of the present study but is an interesting question for future research.

## Conclusions

The results of this study showed that using the surrogate sires strategy could significantly increase the genetic merit of commercial sires, by as much as 6.5 to 9.2 years’ worth of genetic gain, compared to the conventional multiplication strategy. The simulations suggest that identifying elite donors for surrogate sires should be based on three stages, the first of which uses a genomic test followed by two subsequent progeny tests. The use of one or a handful of elite donors to generate surrogate sires that in turn give rise to all production animals would be very different to current practice. While the results demonstrate the great potential of surrogate sires strategy there are considerable risks as well as opportunities. Practical implementation of surrogate sires strategy would need to account for these.

## Authors’ contributions

JMH conceived the study. JMH, MB, and PG designed the study. PG performed the analysis. JMH and PG wrote the first draft of the manuscript. GG, MB, RCG, JJ, RRF, CBAW, AJM, and WOH helped to interpret the results and refine the manuscript. All authors read and approved the final manuscript.

## Acknowledgements

The authors acknowledge the financial support from the BBSRC ISPG to The Roslin Institute BBS/E/D/30002275, from Grant Nos. BB/N015339/1, BB/L020467/1, BB/M009254/1, from Genus PLC and from Innovate UK. This work has made use of the resources provided by the Edinburgh Compute and Data Facility (ECDF) (http://www.ecdf.ed.ac.uk).

## Supplementary material

**Fig. S1.**
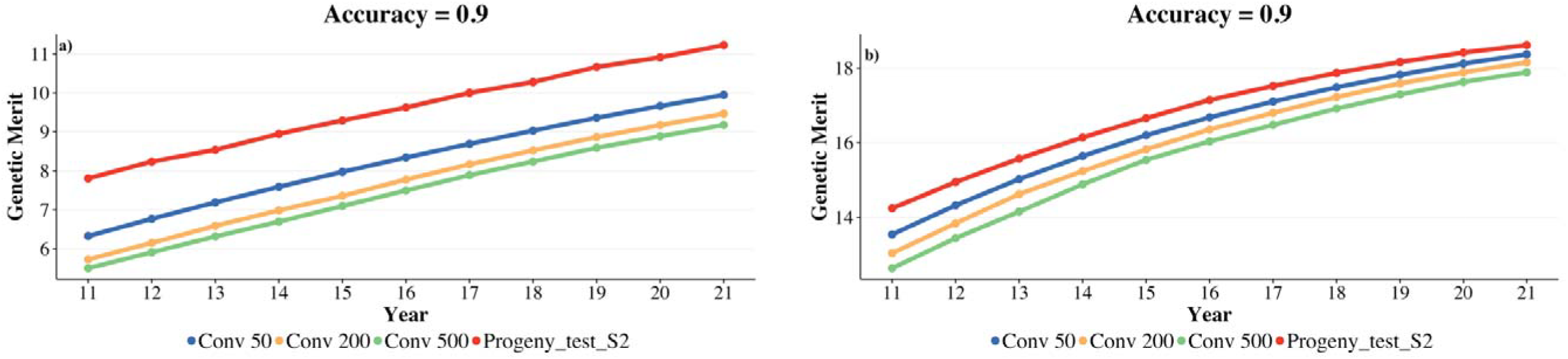
Average genetic merit of commercial sires derived from the best performing surrogate sire strategy scenario and the conventional strategy (top 50, 200 and 500 males) for SmallScenario (a) and BigScenario(b) plotted against time

**Table S1.** Average Years’ worth of Genetic Gain (YGG) with the three-stage testing scenarios of the surrogate sire strategy with one elite donor above the conventional strategy that uses 50 males (BigScenario)

| Progeny test resources<sup>1</sup> | Males progeny tested S1 | Progeny/Male S1 | Males progeny tested S2 | Progeny/Male S2 | YGG<sub>0.5</sub><sup>2</sup> | YGG<sub>0.7</sub><sup>2</sup> | YGG<sub>0.9</sub><sup>2</sup> |
| --- | --- | --- | --- | --- | --- | --- | --- |
| 2000S1/12000S2 | 100 | 20 | 10 | 1200 | 2.2 | 1.5 | 0.9 |
|  |  |  | 20 | 600 | 2.3 | 1.7 | 0.9 |
|  | 200 | 10 | 10 | 1200 | 2.1 | 1.6 | 1.0 |
|  |  |  | 20 | 600 | 2.2 | 1.1 | 0.9 |
|  | 400 | 5 | 10 | 1200 | 2.2 | 1.3 | 0.9 |
|  |  |  | 20 | 600 | 2.3 | 1.5 | 0.8 |
| 4000S1/10000S2 | 100 | 40 | 10 | 1000 | 2.2 | 1.6 | 0.8 |
|  |  |  | 20 | 500 | 2.1 | 1.6 | 0.8 |
|  | 200 | 20 | 10 | 1000 | 2.1 | 2.1 | 0.9 |
|  |  |  | 20 | 500 | 2.2 | 2.1 | 1.0 |
|  | 400 | 10 | 10 | 1000 | 2.3 | 1.7 | 0.9 |
|  |  |  | 20 | 500 | 2.3 | 2.0 | 0.9 |
| 6000S1/8000S2 | 100 | 60 | 10 | 800 | 2.5 | 2.0 | 1.0 |
|  |  |  | 20 | 400 | 2.7 | 2.1 | 1.1 |
|  | 200 | 30 | 10 | 800 | 2.4 | 1.9 | 1.1 |
|  |  |  | 20 | 400 | 2.6 | 2.1 | 1.2 |
|  | 400 | 15 | 10 | 800 | 2.3 | 2.0 | 0.9 |
|  |  |  | 20 | 400 | 2.4 | 2.0 | 0.9 |

<sup>1</sup>Number of total progeny allocated in the first progeny test(S1) and in the second progeny test(S2)
<sup>2</sup>Genomic test accuracy at the initial stage (S0) 0.5, 0.7, and 0.9

**Table S2.** Average Years’ worth of Genetic Gain (YGG) with the three-stage testing scenarios of the surrogate sire strategy with five elite donors above the conventional strategy that uses 50 males (BigScenario)

| Progeny test resources<sup>1</sup> | Males progeny tested S1 | Progeny/Male S1 | Males progeny tested S2 | Progeny/Male S2 | YGG<sub>0.5</sub><sup>2</sup> | YGG<sub>0.7</sub><sup>2</sup> | YGG<sub>0.9</sub><sup>2</sup> |
| --- | --- | --- | --- | --- | --- | --- | --- |
| 2000S1/12000S2 | 100 | 20 | 10 | 1200 | 1.7 | 1.2 | 0.6 |
|  |  |  | 20 | 600 | 1.6 | 1.2 | 0.8 |
|  | 200 | 10 | 10 | 1200 | 1.6 | 1.2 | 0.6 |
|  |  |  | 20 | 600 | 1.5 | 1.0 | 0.3 |
|  | 400 | 5 | 10 | 1200 | 1.6 | 1.1 | 0.7 |
|  |  |  | 20 | 600 | 1.6 | 1.2 | 0.8 |
| 4000S1/10000S2 | 100 | 40 | 10 | 1000 | 1.7 | 1.3 | 0.7 |
|  |  |  | 20 | 500 | 1.7 | 1.2 | 0.6 |
|  | 200 | 20 | 10 | 1000 | 1.6 | 1.3 | 0.4 |
|  |  |  | 20 | 500 | 1.6 | 1.4 | 0.6 |
|  | 400 | 10 | 10 | 1000 | 1.3 | 1.3 | 0.3 |
|  |  |  | 20 | 500 | 1.4 | 1.4 | 0.4 |
| 6000S1/8000S2 | 100 | 60 | 10 | 800 | 1.8 | 1.1 | 0.4 |
|  |  |  | 20 | 400 | 1.8 | 1.4 | 0.7 |
|  | 200 | 30 | 10 | 800 | 1.6 | 1.2 | 0.5 |
|  |  |  | 20 | 400 | 1.6 | 1.2 | 0.5 |
|  | 400 | 15 | 10 | 800 | 1.7 | 1.2 | 0.7 |
|  |  |  | 20 | 400 | 1.6 | 1.2 | 0.9 |

<sup>1</sup>Number of total progeny allocated in the first progeny test(S1) and in the second progeny test(S2)
<sup>2</sup>Genomic test accuracy at the initial stage (S0) 0.5, 0.7, and 0.9

